# Versatile Vasculature Chips for Ultrasound Localization Microscopy

**DOI:** 10.1101/2025.06.15.659747

**Authors:** Renxian Wang, Qi Liu, Xin Zhao, Wei-Ning Lee

## Abstract

Ultrasound localization microscopy (ULM) has revolutionized microvasculature imaging by surpassing the diffraction limit via microbubbles. ULM demonstrates exceptional potential to resolve micrometer-scale vascular structures in both preclinical and clinical studies. However, its performance evaluation remains challenging primarily due to the lack of reference microvascular phantoms featuring micrometer-scale, hierarchical branches, and realistic vascular structures. Inspired by microfluidic chip techniques, we present an organ-on-a-chip protocol for fabricating agarose-based micro-vessel network phantoms with ground truth. The vasculature pattern offers design versatility, enabling on-demand customization. We experimentally demonstrated the feasibility of the vasculature phantom using two adapted patterns. The first was a leaf pattern, which exhibited intrinsic quasi-two-dimensional venation network with hierarchical and branching channels similar to animal vasculature. The second was a kidney pattern, which was based on a two-dimensional projection of real human vasculature obtained from micro computed tomography. The microbubble solution was perfused into the phantoms by capillary force and gravity. The ULM-reconstructed vasculature maps agreed well with the ground truth. ULM achieved a high sensitivity of 0.97 and 0.95, but a low precision of 0.37 and 0.60, for the leaf and kidney phantom, respectively. The results indicated the capability of ULM to reconstruct vessel structures while making many false positive predictions. The proposed protocol holds significant promise for the development and optimization of ultrasound microvascular imaging techniques.

## 1. Introduction

Ultrasound localization microscopy (ULM) is an acoustic super-resolution imaging technique for mapping microcirculation in a living body with the help of gas-filled microbubbles (MBs) that are injected and flow in the circulatory system (Errico *et al*. 2015). Through precise localization and tracking of individual micron-level MBs, ULM overcomes the acoustic diffraction limit and achieves unprecedented micrometric resolution for deep tissue imaging *in vivo*, closing the gap across various imaging modalities (Christensen-Jeffries *et al*. 2020). ULM has shown great potential for broad clinical applicability across various biological tissues, including the brain (Heiles *et al*. 2022, Lowerison *et al*. 2022, Demene *et al*. 2021), kidney (Heiles *et al*. 2022, Chabouh *et al*. 2024), heart (Yan *et al*. 2024), lymph node (Yan *et al*. 2022), and so on. ULM is a multi-step framework, typically consisting of ultrasound data acquisitions of serial frames, MB signal extraction from raw data, MB localization for subpixel accuracy and tracking, and then accumulation of sub-wavelength tracks for final vasculature. Researchers have been working for over ten years on improving the accuracy and efficiency of the ULM workflow, involving various localization and tracking algorithms (Heiles *et al*. 2022, Lerendegui *et al*. 2024, Yan *et al*. 2022, Chen *et al*. 2023, Song *et al*. 2018).

However, validation of ULM remains a critical bottleneck due to the lack of realistic vasculature phantoms that serve as ground truth. The μm diameter, branching, and hierarchical nature of blood vessels makes it challenging to fabricate microvasculature phantoms for assessment of ULM-reconstructed results. The existing validation methods for ULM mainly rely on simulations, manually-assembled tube phantoms with simple geometries, and micro-CT (micro computed tomography). A research group used the Verasonics Research Ultrasound Simulator (Heiles *et al*. 2022) to model each MB as a point scatterer with addictive white Gaussian noise. PROTEUS (Blanken *et al*. 2024) is a recently developed simulator that incorporates flow dynamics for contrast-enhanced ultrasound imaging of realistic vascular structures. It is a customizable framework for ULM validation. A phantom with two crossed tubes of around 200 μm in diameter was commonly used for ULM validation (Zhang *et al*. 2019, Riemer *et al*. 2023, Wang *et al*. 2024). Recently, one study (Parra Raad *et al*. 2024) proposed to manually fabricate a hierarchical microvascular phantom with the diameter down to around λ/6 (~ = ~350 μm) based on polydimethylsiloxane (PDMS). Kawara et al. (Kawara *et al*. 2023) fabricated capillary-scale hydrogel microchannel networks, realizing micrometer scale flow channels but simple branches. Some studies (Andersen *et al*. 2021, Hansen *et al*. 2025) employed micro-CT to validate ultrasound microvascular imaging, but the spatial registration between the two imaging modalities is nontrivial. Overall, simulations can emulate customized and realistic microstructures but hardly replicate actual experimental data; existing phantoms generally do not incorporate micron-level sizes, hierarchical branches, and realistic vascular structures concurrently.

To address the aforementioned challenges, we propose an on-chip versatile vasculature phantom protocol for ultrasound microvascular imaging, inspired by a leaf-templated microfluidic chip technique (Mao *et al*. 2018, Mao *et al*. 2020, Fang *et al*. 2023, Ji *et al*. 2023), with the initial application focused on ULM. We adapted microvasculature patterns as microfluidic channels with sizes of tens of microns and realistic hierarchical branching structures on an agarose layer. Customization of each microvascular structure served as a structural ground truth. In the following sections, we first introduce the protocol and then present experimental results, demonstrating its effectiveness and potential for the development and clinical adoption of ultrasound microvascular imaging techniques.

## 2. Methods

Our protocol follows the typical fabrication workflow of a microfluidic chip, incorporating customized vasculature patterns. As figure 1(a) shows, we binarized the photo of a target pattern provided by the public datasets (Li *et al*. 2022, Walsh *et al*. 2021) and converted it into a vectograph by AutoCAD 2022. The vectograph was then used as a photomask for subsequent photolithography on a silicon wafer to replicate the pattern. A 3% agarose solution was then cast onto the silicon mould. The agarose with the negative pattern of the target vasculature was peeled off from the silicon mould after approximately 10 minutes of solidification at 25 °C.

**Figure 1.**
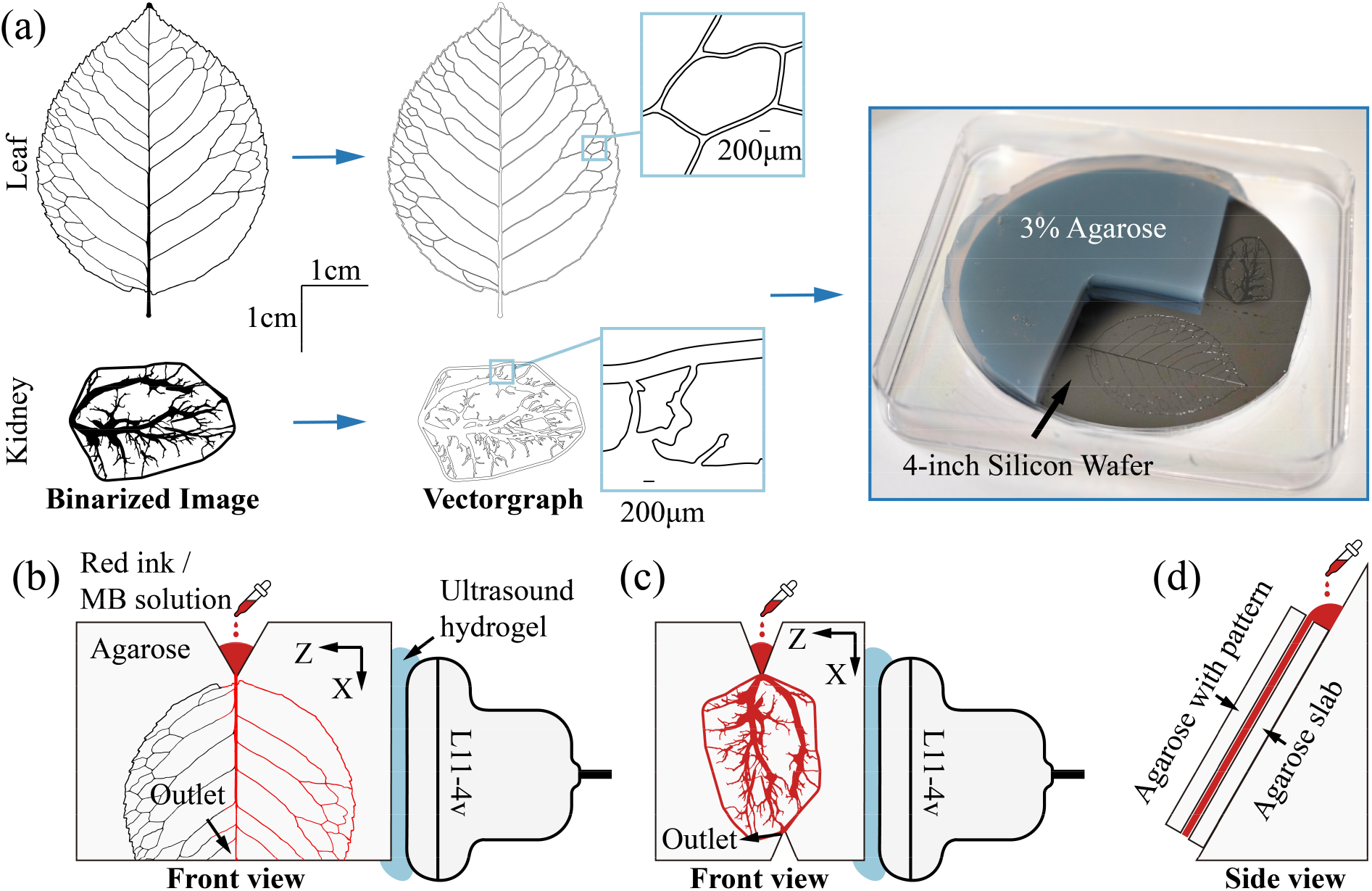
The proposed fabrication protocol of on-chip versatile vasculature phantoms for ultrasound microvascular imaging. (a) Vasculature photos were binarized and converted to vectographs for photolithography on a 4-inch silicon wafer. Micro-channels with the designed vasculature patterns were replicated from the silicon wafer, and the channel depth is approximately 240 μm. (b-d) The ultrasound imaging set-up for data acquisitions. (b) and (c) show the front view of the leaf and kidney phantoms, respectively, and (d) shows the side view. MB denotes microbubble. The red ink is to illustrate perfused vascular regions.

The protocol allows for versatile customization of the vascular network. In this study, we used two vascular patterns (figure 1(a)) adapted from a Chinese rose leaf (Li *et al*. 2022) and a human kidney (Walsh *et al*. 2021). The leaf venation comprised quasi-two-dimensional (2D), hierarchical, and branching channels with high similarity to animal vasculature. As shown in figure 1(b-d), the agarose layer with vasculature network was then tightly sealed to a flat agarose slab (3% agarose, thickness: ~ 6 mm) for MB perfusion and ultrasound imaging. The inlet and outlet were cut out to expose the stem channels of the vasculature network for better perfusion. The MB solution (2×10^7^ to 3×10^7^ MBs/mL, USphere, Taipei) was administered at the inlet and driven by capillary force and gravity.

Ultrasound data were acquired at a compounded frame rate of 400 Hz with 5 tilted plane waves (−4; −2; 0; 2; 4°) using a linear probe L11-4v (6.25 MHz, 128 elements, pitch = 0.30 mm) connected to a Verasonics Vantage 256 system (Verasonics Inc, Kirkland, WA, USA). Each acquisition lasted for four seconds as a bloc, commencing immediately from the start of MB solution perfusion. The total number of blocs was 50 for the leaf phantom and 30 for the kidney phantom.

The PALA ULM workflow was then adopted to reconstruct the vasculature network. It began with MB signal extraction by singular value decomposition. We then employed a Gaussian-fitting method for MB localization (Heiles *et al*. 2022) and a Hungarian method (Kuhn 1955) for MB tracking. The ULM-derived vasculature was lastly compared with the corresponding ground truth obtained from its original AutoCAD design.

We considered the vessel ground truth from the designed pattern in the AutoCAD file as a binarized mask, with the vascular network region as the target. For quantitative comparison, ULM results were binarized using different thresholds (*th*); pixels with intensities greater than or equal to *th* were assigned a value of 1 (positive), while all other pixels were assigned a value of 0 (negative). The ULM results were manually registered to ground truth. Positive pixels in the binarized ULM results that fell within the vascular region of the ground truth were regarded as true positives (TPs), while those in the non-vascular region were considered false positives (FPs). Negative pixels in the binarized ULM results that actually corresponded to the vascular region of the ground truth were labelled as false negatives (FNs). Precision, sensitivity, and the Jaccard (JAC) index were computed for quantitative evaluation of the ULM results and respectively defined as

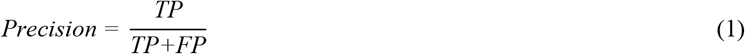

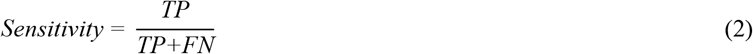

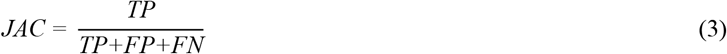

## 3. Results

Figure 2(a) shows the ULM vasculature map of the leaf phantom, corresponding to a 90° counterclockwise rotation of the imaging view shown in figure 1(b). The upper part of the leaf venation network was reconstructed well, but the rest was not. The light blue box in figure 2(a) was zoomed-in for better visualization (figure 2(b)) and binarized (figure 2(c)) with *th* = 0 for comparison with the ground truth (figure 2(d)). Figure 2(e) shows the cross-sectional vessel profiles along lines 1, 2, and 3 in figure 2(b) with measured diameters (69.0 μm, 51.5 μm, and 58.0 μm, respectively) around or below λ / 4 (λ = 246 μm). The ULM-estimated diameters were about 45-75% of the ground truth values. The peaks of the ULM intensity profiles were all located within the ground truth vessels.

**Figure 2.**
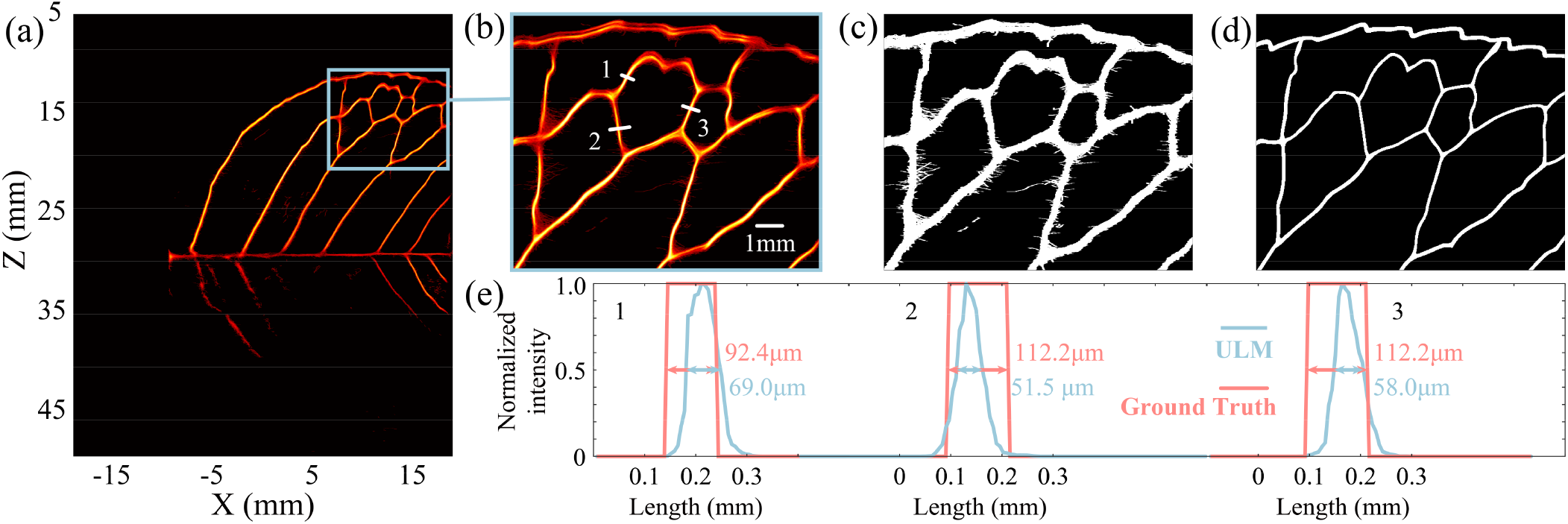
ULM of a leaf phantom: (a) The ULM density map of the leaf phantom in the front view (90° counterclockwise rotation of figure 1(b)). (b) Magnification of the venation network enclosed by the light blue box in (a). (c) Binarized vasculature map of (b) with *th* = 0. (d) The corresponding vasculature ground truth for (b) and (c), where (b-d) share the same scale bar. (e) The intensity profiles along the three white lines in (b). The light blue and red lines denote the ULM results and the ground truths, respectively.

The kidney pattern was much more challenging to reconstruct because it has more hierarchical and branching structures. Yet, its ULM result (figure 3(a)) was in good qualitative agreement with the ground truth (figure 3(c)) despite some deviations in their details. The main branches were delineated well in the ULM map, while some complex microchannels were not reconstructed, particularly the cul-de-sac channels indicated by turquoise triangles (figure 3). ULM also did not detect vessels in the positions of purple triangles (figure 3). Some MBs flowed outside the micro-channels, and their flow paths were delineated as free trajectories (orange triangles in figure 3).

**Figure 3.**
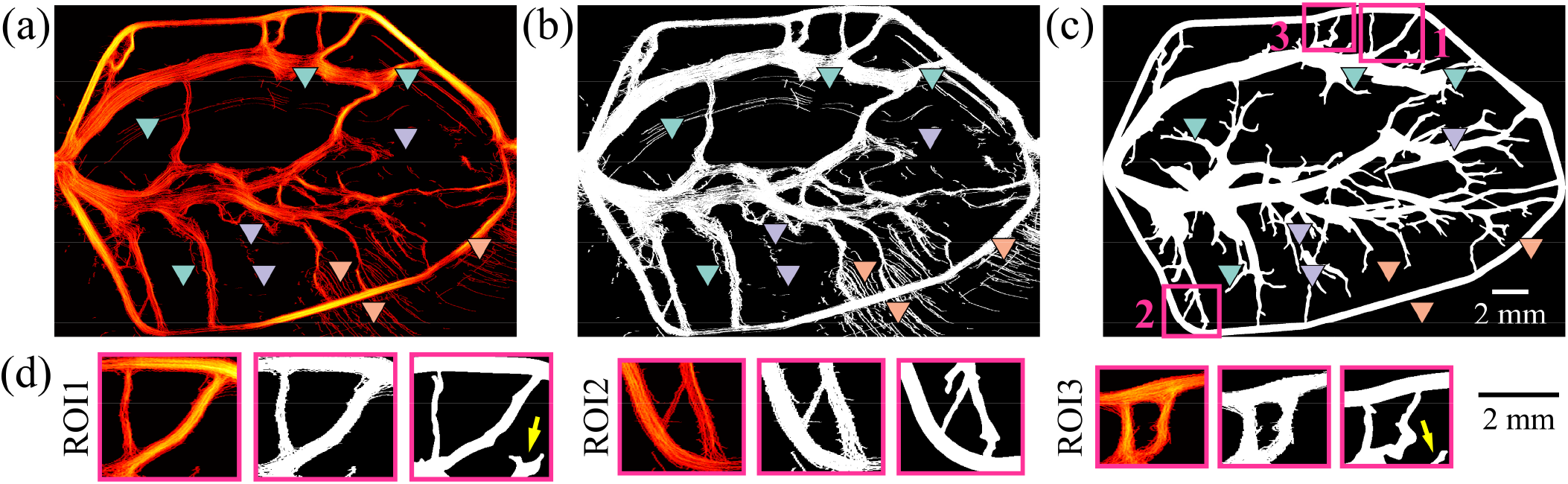
ULM of a kidney phantom: (a) The ULM density map of the kidney phantom. (b) Binarized vasculature map of (a) with the threshold (*th*) of 0. (c) The corresponding vasculature ground truth. (d) Side-by-side comparison of three zoomed-in ROIs enclosed by the magenta boxes in (c) across maps in (a-c). Note that (a-c) share the same scale bar.

Figure 4(a) shows the quantitative analysis of the leaf ULM map enclosed by the light blue box (figure 2(a)), which exhibited complex network. Empirical thresholds (0, 0.01, 0.025, 0.05, 0.1, 0.2, 0.3) were applied to the leaf ULM result for binarization and subsequent evaluation. As the threshold increased, the precision increased from 0.37 to 0.84 while the sensitivity decreased from 0.97 to 0.27. The JAC index reached its maximum, approximately 0.51, at the threshold of 0.01. Figure 4(b) shows the quantitative analysis of the ULM result for the kidney phantom from the average of the three ROIs with branching structures in figure 3(d). The three ROIs corresponded to the three locations labelled in figure 3(c). The thresholds were selected empirically as 0, 0.005, 0.01, 0.02, 0.035, 0.05, and 0.1 for binarization on the three ROIs. As the threshold increased, the precision increased from 0.60 to 0.92, while the sensitivity dropped from 0.95 to 0.37. The JAC index initially increased to the maximum, approximately 0.70, when *th* = 0.005, and then decreased to 0.35 at *th* = 0.1. Note that the aforementioned cul-de-sac regions were not included in the quantitative analysis of the three ROIs (marked by the yellow arrows in figure 3(d)).

**Figure 4.**
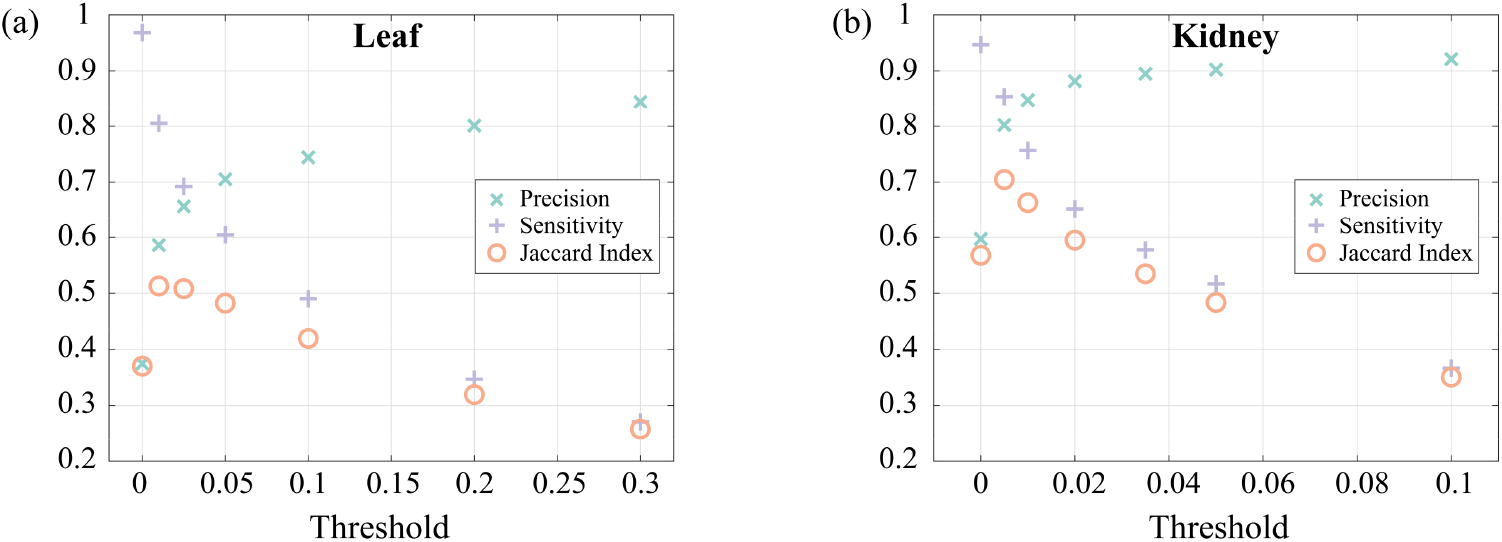
Quantitative analysis of ULM at different binarization thresholds for (a) the leaf and (b) the kidney phantoms. (a) and (b) correspond to the ROIs in figure 2(b) and figure 3(d), respectively.

## 4. Discussion

We introduced an on-chip versatile vasculature phantom protocol that enables flexible customization of micron-sized and hierarchical microvasculature. The phantom medium is agarose, which is both acoustically and optically transparent. Optical imaging provided more flow information for further assessment (supplementary videos 1-3), and the designed pattern in the AutoCAD file served as a structural ground truth. The minimum diameter of the fabricated channels was approximately 50 μm, far below the acoustic diffraction limit of most clinical and research-used ultrasonic settings. The fabricated phantom can validate not only ULM but also other ultrasound and non-ultrasound microvascular imaging techniques.

ULM resolution was assessed using the leaf phantom, showing that ULM-estimated diameters of the microchannels (figure 2(e)) at half-maximum intensity were smaller than the ground truth values. This was because ULM relied on MB flow dynamics, and the MB signals extracted by SVD was related to the velocity distribution within the flow channel, resulting in a quasi-parabolic intensity profile (figure 2(e)). For both leaf and kidney phantoms, the high sensitivity (0.97 and 0.95) but low precision (0.37 and 0.60) at *th* = 0 indicated that ULM successfully reconstructed the vessels within the selected ROIs but with a large number of FPs, particularly in the branching regions (figure 2(c) and 3(d)). As the binarization threshold increased, precision and JAC index increased while sensitivity decreased in both phantom cases. Yet, the kidney case had a higher precision than the leaf one. We attributed this observation to the less complex microvasculature in the selected ROIs of the kidney case than that in the leaf case, which contained many branches. This finding highlighted the challenge for ULM to reconstruct branching structures more complex than straight channels.

ULM could not reconstruct the lower half of the leaf vasculature in the deep image zone probably due to channel congestion by air. This could be caused by the pumpless nature of our phantom, and the MB solution was perfused by capillary force and gravity. The kidney phantom had a more severe problem with air congestion (purple triangles in figure 3 and supplementary video 2) because of its much more complex vasculature, including multiscale microvascular network and more bifurcations, than that in the leaf phantom. The kidney vasculature map reconstructed by ULM generally agreed with the ground truth image, but its quality was multifactorial. The cul-de-sac channels, marked by the turquoise triangles in figure 3 and shown in the supplementary video 1, had no circulatory flow so that ULM failed to detect them. Since the agarose microfluidic chip was sealed spontaneously by hydrogen bonds, the adhesion could not be guaranteed as reliably as that of traditional PDMS-based microfluidic chips, especially at high flow velocities. In our experiments, fluid leakage occurred in the areas that were indicated by the orange triangles in figure 3 and supplementary video 3, where free MB trajectories were delineated by ULM. It is important to note that the issue of fluid leakage could be mitigated through proper pattern design, such as avoiding cul-de-sac channels. This may explain why there was no fluid leakage observed in the leaf phantom.

Manual spatial registration between ULM and ground truth images may have led to alignment errors although the high sensitivity at *th* = 0 indicated good registration. In particular, the channel depth of our phantoms was 240 μm, which was deemed to be relatively large for a typical microfluidic chip. The large channel depth was designed to facilitate pumpless perfusion but might cause photolithography defect and could exacerbate the mis-alignment between the ultrasound imaging plane and the vasculature channel. Moreover, generation of ULM images involved many parameters in the localization and tracking steps, including SVD cut-off threshold, the number of MBs per frame, localization methods, the minimum trajectory length, and so on. Finding the optimal set of parameters is an exhaustive task. The parameters that were used for the leaf and kidney data in this study were summarized in Table S1 of the Supplementary Material.

Overall, the proposed on-chip vasculature protocol provides a valuable method to validate and assess the performance of ULM, showing significant potential for the development and optimization of ultrasound microvascular imaging techniques. The methodology also paves the way for developing versatile organ-on-a-chip devices. Future work could focus on performance evaluation of volumetric ULM and micro-Doppler imaging with flow velocity information.

## Data availability statements

The data that support the findings of this study are openly available at the following URL/DOI: https://doi.org/10.5281/zenodo.15614916. Data will be available from 01 December 2025.

## Acknowledgements

Renxian Wang is supported by ACCESS – AI Chip Center for Emerging Smart Systems, sponsored by the InnoHK initiative of the Innovation and Technology Commission of the Hong Kong Special Administrative Region Government. This work was in part supported by Collaborative Research with World-leading Research Groups from The Hong Kong Polytechnic University (P0039523).

## References

Andersen S B, Taghavi I, Kjer H M, Sogaard S B, Gundlach C, Dahl V A, Nielsen M B, Dahl A B, Jensen J A and Sorensen C M 2021 Evaluation of 2D super-resolution ultrasound imaging of the rat renal vasculature using ex vivo micro-computed tomography Sci Rep 11 24335

Blanken N, Heiles B, Kuliesh A, Versuis M, Jain K, Maresca D and Lajoinie G 2024 PROTEUS: A Physically Realistic Contrast-Enhanced Ultrasound Simulator-Part I: Numerical Methods IEEE Trans Ultrason Ferroelectr Freq Control PP

Chabouh G, Denis L, Bodard S, Lager F, Renault G, Chavignon A and Couture O 2024 Whole Organ Volumetric Sensing Ultrasound Localization Microscopy for Characterization of Kidney Structure IEEE Trans Med Imaging 43 4055–63

Chen X, Lowerison M R, Dong Z, Chandra Sekaran N V, Llano D A and Song P 2023 Localization Free Super-Resolution Microbubble Velocimetry Using a Long Short-Term Memory Neural Network IEEE Trans Med Imaging 42 2374–85

Christensen-Jeffries K, Couture O, Dayton P A, Eldar Y C, Hynynen K, Kiessling F, O’Reilly M, Pinton G F, Schmitz G, Tang M X, Tanter M and van Sloun R J G 2020 Super-resolution Ultrasound Imaging Ultrasound Med Biol 46 865–91

Demene C, Robin J, Dizeux A, Heiles B, Pernot M, Tanter M and Perren F 2021 Transcranial ultrafast ultrasound localization microscopy of brain vasculature in patients Nat Biomed Eng 5 219–28

Errico C, Pierre J, Pezet S, Desailly Y, Lenkei Z, Couture O and Tanter M 2015 Ultrafast ultrasound localization microscopy for deep super-resolution vascular imaging Nature 527 499–502

Fang J, Liu H, Qiao W, Xu T, Yang Y, Xie H, Lam C H, Yeung K W K and Zhao X 2023 Biomimicking Leaf-Vein Engraved Soft and Elastic Membrane Promotes Vascular Reconstruction Adv Healthc Mater 12 e2201220

Hansen L N, Ráth A, Amin Naji M, McDermott A, Sørensen C M, Kjer H M, Gundlach C, Dahl A B and Jensen J A 2025 Assessment of Super-Resolution Ultrasound Imaging Using the Erythrocytes Through Comparison with Micro-CT

Heiles B, Chavignon A, Hingot V, Lopez P, Teston E and Couture O 2022 Performance benchmarking of microbubble-localization algorithms for ultrasound localization microscopy Nat Biomed Eng 6 605–16

Ji X, Bei H P, Zhong G, Shao H, He X, Qian X, Zhang Y and Zhao X 2023 Premetastatic Niche Mimicking Bone-On-A-Chip: A Microfluidic Platform to Study Bone Metastasis in Cancer Patients Small 19

Kawara S, Cunningham B, Bezer J, Kc N, Zhu J, Tang M X, Ishihara J, Choi J J and Au S H 2023 Capillary-Scale Hydrogel Microchannel Networks by Wire Templating Small 19 1–10

Kuhn H W 1955 The Hungarian method for the assignment problem Naval Research Logistics Quarterly 2 83–97

Lerendegui M, Riemer K, Papageorgiou G, Wang B, Arthur L, Chavignon A, Zhang T, Couture O, Huang P, Ashikuzzaman M, Dencks S, Dunsby C, Helfield B, Jensen J A, Lisson T, Lowerison M R, Rivaz H, Samir A E, Schmitz G, Schoen S, van Sloun R, Song P, Stevens T, Yan J, Sboros V and Tang M X 2024 ULTRA-SR Challenge: Assessment of Ultrasound Localization and TRacking Algorithms for Super-Resolution Imaging IEEE Trans Med Imaging 43 2970–87

Li L, Hu W, Lu J and Zhang C 2022 Leaf vein segmentation with self-supervision Computers and Electronics in Agriculture 203

Lowerison M R, Sekaran N V C, Zhang W, Dong Z, Chen X, Llano D A and Song P 2022 Aging-related cerebral microvascular changes visualized using ultrasound localization microscopy in the living mouse Sci Rep 12 619

Mao M, Bei H P, Lam C H, Chen P, Wang S, Chen Y, He J and Zhao X 2020 Human-on-Leaf-Chip: A Biomimetic Vascular System Integrated with Chamber-Specific Organs Small 16 e2000546

Mao M, He J, Lu Y, Li X, Li T, Zhou W and Li D 2018 Leaf-templated, microwell-integrated microfluidic chips for high-throughput cell experiments Biofabrication 10 025008

Parra Raad J, Lock D, Liu Y Y, Solomon M, Peralta L and Christensen-Jeffries K 2024 Optically Validated Microvascular Phantom for Super-Resolution Ultrasound Imaging IEEE Trans Ultrason Ferroelectr Freq Control 71 1833–43

Riemer K, Toulemonde M, Yan J, Lerendegui M, Stride E, Weinberg P D, Dunsby C and Tang M X 2023 Fast and Selective Super-Resolution Ultrasound In Vivo With Acoustically Activated Nanodroplets IEEE Trans Med Imaging 42 1056–67

Song P, Trzasko J D, Manduca A, Huang R, Kadirvel R, Kallmes D F and Chen S 2018 Improved Super-Resolution Ultrasound Microvessel Imaging With Spatiotemporal Nonlocal Means Filtering and Bipartite Graph-Based Microbubble Tracking IEEE Trans Ultrason Ferroelectr Freq Control 65 149–67

Walsh C L, Tafforeau P, Wagner W L, Jafree D J, Bellier A, Werlein C, Kuhnel M P, Boller E, Walker-Samuel S, Robertus J L, Long D A, Jacob J, Marussi S, Brown E, Holroyd N, Jonigk D D, Ackermann M and Lee P D 2021 Imaging intact human organs with local resolution of cellular structures using hierarchical phase-contrast tomography Nat Methods 18 1532–41

Wang B, Riemer K, Toulemonde M, Yan J, Zhou X, Smith C A B and Tang M X 2024 Broad Elevation Projection Super-Resolution Ultrasound (BEP-SRUS) Imaging With a 1-D Unfocused Linear Array IEEE Trans Ultrason Ferroelectr Freq Control 71 255–65

Yan J, Huang B, Tonko J, Toulemonde M, Hansen-Shearer J, Tan Q, Riemer K, Ntagiantas K, Chowdhury R A, Lambiase P D, Senior R and Tang M X 2024 Transthoracic ultrasound localization microscopy of myocardial vasculature in patients Nat Biomed Eng 8 689–700

Yan J, Zhang T, Broughton-Venner J, Huang P and Tang M X 2022 Super-Resolution Ultrasound Through Sparsity-Based Deconvolution and Multi-Feature Tracking IEEE Trans Med Imaging 41 1938–47

Zhang G, Harput S, Hu H, Christensen-Jeffries K, Zhu J, Brown J, Leow C H, Eckersley R J, Dunsby C and Tang M X 2019 Fast Acoustic Wave Sparsely Activated Localization Microscopy (fast-AWSALM): Ultrasound Super-Resolution using Plane-Wave Activation of Nanodroplets IEEE Trans Ultrason Ferroelectr Freq Control

